# Towards operational epidemiology of *Pyricularia oryzae* in sports turf: contribution of a multi-matrix molecular monitoring strategy at stadium scale

**DOI:** 10.64898/2026.09.30.755619

**Authors:** Saadi Sarah, Gruet Cécile, Adreit Henri, Poncelet Nicolas, Di Mattia Jérémy, Courot Olivier, Tharreau Didier

**Affiliations:** CIRAD, UMR PHIM, Montpellier, France; PHIM, Univ. Montpellier, CIRAD, INRAE, IRD, Institut Agro Montpellier, Montpellier, France; IAGE

**Keywords:** *Pyricularia oryzae*, Gray leaf spot, molecular biology, digital PCR, early detection, surveillance, disease monitoring, stadium, turfgrass

## Abstract

Gray leaf spot can cause severe turfgrass damage, particularly in intensively managed sports fields, while disease management remains challenging due to the rapid development of symptoms under favorable environmental conditions. Early detection of *Pyricularia oryzae* could therefore provide valuable information for anticipating disease development and supporting management decisions. This study aimed to develop a digital PCR (dPCR) molecular tool to support the monitoring and management of gray leaf spot disease in stadium turfgrass. The assay showed high sensitivity, consistently detecting *P. oryzae* when infected tissue represented only 0.05% of a mixture with healthy turfgrass, and enabled pathogen detection before symptom appearance under both controlled and field conditions. Different matrices were also investigated across 15 stadiums to identify the most relevant material for disease monitoring. Roots showed the highest prevalence of *P. oryzae* detection, including during asymptomatic periods, highlighting their potential as a relevant matrix for pathogen surveillance. In a stadium monitored several months after a gray leaf spot outbreak, *P. oryzae* DNA was still detected in roots from 27 of the 32 sampled zones despite the absence of visible symptoms. Overall, these results demonstrate the potential of dPCR for the early detection and monitoring of *P. oryzae* in stadium turfgrass, particularly through root sampling during periods when environmental conditions are not favorable for gray leaf spot development, and a spatiotemporal surveillance at stadium scale.

## 1. Introduction

Fungal pathogens represent a major threat to food crops, ornamentals, and plants for recreative areas. Among them, *Pyricularia oryzae* Cavara (synonym *Magnaporthe oryzae*) is one of the most destructive fungal pathogens affecting *Poaceae* worldwide (Viji et al. 2001). This hemibiotrophic ascomycete infects at least 20 grass species, including major crops such as rice, wheat, millet, maize, and numerous turfgrass species (Couch et al. 2005, Gladieux et al. 2018, Pordel et al. 2021, Baudin et al. 2024). The pathogen is well known as the causal agent of rice blast disease, and of gray leaf spot (GLS) disease in turfgrass systems, particularly on perennial ryegrass (*Lolium perenne*) (Tosa et al. 2007).

GLS has become a major disease of sport’s turf over the last three decades. First reported on perennial ryegrass turf in Pennsylvania in 1992, it subsequently caused severe epidemics throughout the United States, particularly on golf courses and athletic fields (Viji et al. 2001, Farman et al. 2002). The first epidemics on turf grass in Japan were reported in 1996 (Tanaka et al. 2000). Since then, GLS has been reported on an increasing number of turfgrass hosts, including tall fescue, St. Augustinegrass, and hard fescue, highlighting the large host range of *P. oryzae*. (Tredway et al. 2005; Vines et al. 2022). In Europe, GLS has emerged more recently, but is now considered a growing threat. The disease was officially reported in France in 2016 on perennial ryegrass sports fields hosting professional football events (Milazzo et al. 2019). The disease results in rapid turf decline and extensive economic losses, with entire turf stands destroyed within only a few days following symptom appearance (Uddin et al. 2003a). The epidemiological success of *P. oryzae* is partly linked to its biological characteristics. The fungus survives as mycelium on infected plant debris and produces abundant airborne conidia under warm and humid conditions (Vincelli 2000). These spores are dispersed by wind, irrigation, splashing water, or machinery, enabling rapid spread in highly managed turf ecosystems. Symptoms may appear only a few days after infection, and multiple infection cycles can occur during a single growing season, leading to explosive outbreaks (Bonos et al. 2006, Huang et al. 2020).

Management of GLS in stadium turf is challenging because stadiums are artificial ecosystems requiring fine-tune management, with a specific microclimate. Sport’s turf is subjected to mechanical stress, intensive irrigation promoting prolonged leaf wetness, high nitrogen fertilization, mowing, and frequent overseeding, all of which can favor GLS development and pathogen spread. Current management mainly relies on cultural practices, resistant cultivars or species, and fungicide applications. However, fungicide efficacy strongly depends on application timing, and curative treatments are often limited once symptoms appear (Williams et al. 2001, Uddin et al. 2003ac, Villari et al. 2017). Fungicides also become ineffective after selection of resistant mutants in populations of *P. oryzae* (Vincelli and Dixon 2002, Kim et al. 2003). Furthermore, recent regulations such as the French “Loi Labbé” have restricted the use of synthetic pesticides in public sports facilities, increasing the need for integrated management approaches combining cultural practices, resistant cultivars (which are not so common in perennial ryegrass), biocontrol products, and optimized disease monitoring systems for early intervention.

Despite the importance of early intervention, current disease diagnosis in sports turf remains largely symptom-based. GLS symptoms can easily be confused with those caused by other summer turf diseases or abiotic stresses, making accurate field diagnosis difficult. Furthermore, visual symptoms generally appear only after substantial pathogen colonization has already occurred, reducing the efficiency of subsequent management interventions. Conventional diagnosis relies on microscopic observation, and fungal isolation from symptomatic tissues (Uddin et al. 2003b) which require several days. To overcome these limitations, several molecular diagnostic approaches have been developed for turfgrass pathogens, including PCR, qPCR, Immuno-recognition assay, and LAMP-based assays (Harmon et al. 2003, Uddin et al. 2003b, Villari et al. 2017, Li et al. 2019, Thierry et al. 2020a). These approaches significantly improved the sensitivity and specificity of pathogen detection and demonstrated the possibility of identifying *P. oryzae* before severe symptom expression. For example, the airborne inoculum of *P. oryzae* was successfully detected up to 12 days before symptom appearance in turfgrass plots using a qLAMP-spore trap system (Villari et al. 2017). Similar molecular approaches have also been developed for other turfgrass fungal pathogens such as *Clarireedia* spp., where pathogen DNA was detected in asymptomatic plant tissues before disease outbreaks (Groben et al. 2020).

Nevertheless, most current diagnostic approaches focus on single matrices, mainly symptomatic foliar tissues, and are designed for laboratory confirmation rather than operational field surveillance. In addition, little is known about the spatial and temporal dynamics of *P. oryzae* within stadium ecosystems and important questions remain regarding inoculum sources, pathogen circulation, persistence, and asymptomatic colonization. Recent advances in multi-matrix molecular monitoring by digital PCR open the way to develop rapid and sensitive tools for GLS diagnostic in different compartments to support integrated and early disease management and to produce epidemiological knowledge.

The objective of this study was to evaluate the contribution of a multi-matrix molecular monitoring strategy based on digital-PCR (dPCR) for GLS management in sport’s turf systems. More specifically, this work aimed to assess the ability of early molecular detection to identify *P. oryzae* before symptom appearance, investigate pathogen distribution across stadium and across matrices, and explore how these data may provide epidemiological insights useful for integrated disease management at stadium scale.

## 2. Materials and methods

### 2.1. Design of specific primers for dPCR detection of *P. oryzae*

For the development of a detection system specific to *P. oryzae*, a total of 60 strains were used. Forty-eight strains of *P. oryzae* were selected from various hosts and genetic lineages to cover the species’ genetic diversity and 12 strains belonging to other *Pyricularia* and *Proxipyricularia* species were used as outgroups to assess specificity (Table S1). The genome sequences of these strains were obtained from the Plant Health Institute of Montpellier (PHIM) unit at the CIRAD Institute, Montpellier, France and from the Nucleotide database of the National Center for Biotechnology Information (NCBI). Five genomic regions (Internal transcribed region (ITS), Large subunit ribosomal RNA gene (LSU), RNA polymerase II subunit (RPB1), Calmodulin gene (CaM), and Actin gene (ACT) commonly used in phylogenetic studies were extracted from the genomes of these strains. Multiple sequence alignments were performed using Multalign, a progressive multiple-alignment algorithm based on hierarchical clustering (Corpet 1988), to identify the gene providing the highest specificity for *P. oryzae*. The gene was selected based on its level of identity between the 48 strains of the species selected and the polymorphism relative to the 12 strains of other species (Table S1). A primer pair and a TaqMan probe were developed using Primer3plus. The specificity of the primers and probe developed was finally verified on the NCBI database.

### 2.2. Validation of specificity and inclusivity of the detection system on genomic DNA of fungal strains

To assess the ability of the designed primers and probe to specifically target *P*. *oryzae*, a validation panel was created. The validation panel was designed to assess both exclusivity and inclusivity by comprising (i) fungal pathogens commonly associated with turfgrass and species phylogenetically related to *Pyricularia* to evaluate analytical specificity, and (ii) a broad collection of *P. oryzae* isolates representing the known diversity of the species, including the extensively characterized rice lineages, to ensure robust detection across the target population. Nineteen *P. oryzae* isolates from different *Poaceae* species, including eleven originating from turfgrass, were used. The study also included 29 isolates representing different species within the genera *Pyricularia*, *Pseudopyricularia* and *Proxipyricularia*, as well as seven isolates of another fungal species pathogenic to turfgrass (Table S1). All isolates originated from the CIRAD fungal collection at the Plant Health Institute of Montpellier (PHIM). Genomic DNA from fungal mycelium was extracted using a CTAB/phenol–chloroform method (Gladieux et al. 2018) and subsequently analyzed by dPCR with the developed markers.

Digital-PCR amplification was conducted on a QIAcuity® Eight Plateform System (QIAGEN, Germany) following manufacturer’s instructions, using the QIAcuity® Probe PCR Kit (Cat. No. 250103) and QIAcuity® Nanoplate 26 K 24-wells (Cat. No. 250001). The dPCR reaction mixtures were prepared in a preplate as follows. For each reaction, 10 µL of QIAcuity® Probe PCR Kit, 1 µL of primers-probe mix, and 2 µL of extracted DNA were combined with H2O up to 40 µL final reaction volume. Reaction mixtures were transferred into a QIAcuity® Nanoplate, and the plate was loaded onto the QIAcuity® Eight instrument, a fully automated system. The workflow included (i) priming and rolling step to generate and isolate the chamber partitions (26,000 partitions), (ii) amplification step with the following cycling protocol: 95°C for 2 min for the enzyme activation, followed by 45 cycles of 20 sec at 95°C for denaturation, and 60 sec of annealing/extension at 60°C, (iii) imaging step by reading fluorescence emission after excitation of the probe at the appropriate wavelength. Data were analyzed using the QIAcuity® Software suite V2.0.20 and expressed as copies per microliter of reaction volume (40 µL).

### 2.3. Temporal survey of perennial ryegrass colonization after infection with *P. oryzae* spores

The kinetics of *P. oryzae* infection was investigated by quantifying fungal DNA of *P. oryzae*, using dPCR, on perennial ryegrass inoculated with spores. A mix blend of three varieties was sown: New Orleans 15%, Colosseum 40%, Ultra 45%. The plants were grown on a 30 x 50 cm culture tray in a greenhouse. After 10 days, once the turf was sufficiently established, the strain FR1084 of *P. oryzae* was inoculated using 20 mL of a spore suspension (25,000 spores.mL-1) prepared in water with gelatin (0.5%). Approximately 0.5 g of samples were collected at 6 time-points: prior to inoculation, immediately after inoculation and at 3, 5, 7, and 10 days post-inoculation. Subsequently, the experiment was repeated to add earlier time-points (prior to inoculation, 1h, 5h, 24h, 48h, 72h, 140h post-inoculation). Three biological replicates were performed for each sampling time. Samples were immediately frozen at –80°C following collection. Samples were then put in a 2 mL tube with 2 beads (4 mm) and ground in liquid nitrogen using a bead mill grinder (GENO/GRINDER2100®, SPEX SamplePrep, Metuchen, USA) for 3 cycles of 20 seconds at 1500 cpm. Then, 0.1 g of grounded material was used for DNA extraction using the Nucleomag plant kit (Ref 744400.4, Macherey-Nagel, Düren, Germany) following manufacturer’s instructions with two modifications (i) the volume of magnetic bead was reduced to 10 µL and (ii) the lysis time was set to 10 min. All extracted samples were subsequently analysed by dPCR according to the protocol described in section 2.2. DNA quantification values were log10-transformed before statistical analysis. Differences among sampling times were evaluated using one-way analysis of variance (ANOVA), after checking assumptions of residual normality using the Shapiro–Wilk test and homogeneity of variances using Levene’s test. Significant ANOVA results were followed by Tukey’s honestly significant difference (HSD) post-hoc test for pairwise comparisons.

### 2.4. Temporal survey of inoculum in a stadium

Aerial parts of turf were collected in three areas of a football stadium from 7 Novembre 2022 to 31 July 2023, once a month until mid-May and then every two weeks. Part of the sample was used for detection of *P. oryzae* by digital PCR, following the method described above (2.3). The remaining part of the sample was used for microbiological observation. Aerial parts of 5 plants were cut and deposited on a filter paper in a Petri dish. The paper was moistened with water, and plates were placed at 25°C to allow sporulation of fungi. Samples were observed every two days for one week under a stereomicroscope and the number of lesions with typical spores of *P. oryzae* was scored for each sample.

### 2.5. Evaluation of the detection limit

The detection limit of the method, including pre-treatment, DNA extraction and analysis, was evaluated using healthy perennial ryegrass mixed with different quantities of *P. oryzae*-infected perennial ryegrass (variety Etienna), collected 140 hours post-inoculation (performed as described above in section 2.3). Healthy and infected grass were ground separately in liquid nitrogen using a knife mill (Pulverisette 11, Fritsch, Idar-Oberstein, Germany; 1 cycle of 10 s at speed of 10 000 rpm). The samples were then pooled and homogenized following 5 different proportions of infected tissue: 0% (negative control), 0.025% (w/w), 0.05%, 0.1%, and 100% (positive control), each in a final sample mass of 50 g of grass. Three independent biological replicates were systematically performed for each condition. Samples were then subjected to DNA extraction and analysed by dPCR using the protocol described above (2.2; 2.3).

### 2.6. Prevalence of *P. oryzae* in stadiums

To investigate the reservoir of *P. oryzae* in stadiums, soil, turf roots, and aerial parts were analyzed in 15 stadiums in France during spring, *i.e.* before the favorable periods for disease development. Sample collection was performed in each stadium by stadium’s management teams, and samples were sent to I.A.G.E for analysis. Grass samples were treated as described before (2.3; 2.5). Roots samples were pretreated by grinding in liquid nitrogen with a knife mill (1 cycle of 10 s at speed of 10 000 rpm) and 0.1 g of material was used for DNA extraction. Soil samples were dried in a vacuum oven (VO101, Memmert, Schwabach, Germany) for 1h30 at 70°C. The dried samples were then placed in 500 mL inox bottles with 2 ceramic grinding cylinder (9.5 × 22.2 mm, SPEX SamplePrep, USA) and homogenized using a bead mill homogenizer (GENO/GRINDER2100®, SPEX SamplePrep, Metuchen, USA) for 3 cycles of 20 seconds at 1500 cpm. For both root and soil samples, a sub-sample (0.1 g for roots and 0.2 g for soil) was then placed in a Macherey-Nagel bead tube A (Ref 740786.50, Macherey-Nagel, Düren, Germany) and 800 µL of MagAttract Power Soil kit (Cat no. 47119, Qiagen, Venlo, The Netherlands) lysis buffer were added to the tubes. Lysis was performed using a bead mill homogenizer for 10 min at 1500 cpm. The purification and washing steps were then carried out according to the kit’s protocol. DNA extracts were then analyzed by dPCR following the protocol described in section 2.2. Differences in *P. oryzae* DNA quantification among the three matrices (soil, roots and aerial parts) were assessed using a Friedman test, considering the paired structure of the data (the three matrices were collected from the same stadiums). When significant differences were detected, pairwise comparisons were performed using Wilcoxon signed-rank tests with Holm correction for multiple comparisons. In order to determine if fungal abundance is correlated between matrices, correlation of DNA quantities in the different matrices was tested using Spearman’s rank correlation coefficient.

### 2.7. Characterization of *P. oryzae* spatial distribution in a stadium

*P. oryzae* spatial distribution was investigated using a stadium recovering from a gray leaf spot outbreak. The stadium developed disease symptoms during the summer, and sampling was done several months later, once disease management practices had been implemented and visible symptoms had disappeared. The field was divided into 32 distinct zones (Figure 1) ensuring uniform and representative coverage of the entire area. Grass and root samples were collected in each zone by the field management. Fifteen randomly distributed individual samples were collected per zone using a turf plugger. Aerial parts were then separated using scissors. The 15 samples from each zone were pooled into a single composite sample per zone, thereby reducing bias associated with spatial variability. Samples were treated as described above (section 2.6).

**Figure 1.**
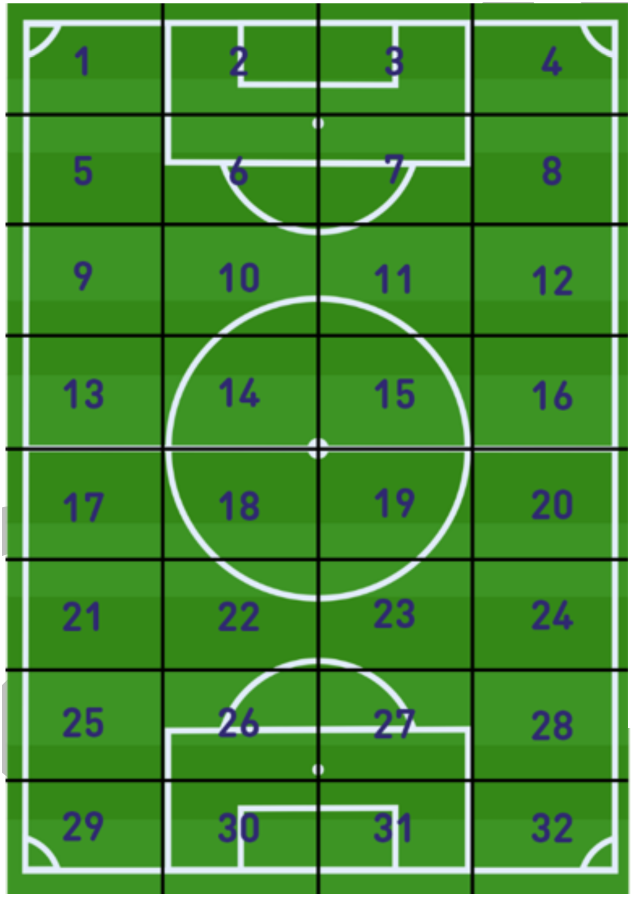
Schematic representation of the 32 distinct sampling areas on the stadium.

## 3. Results

### 3.1. The dPCR assay detects *P. oryzae* with very limited cross-reactivity

Primers and probe were developed on the RPB1 gene. Among the five candidate genes evaluated, RPB1 displayed the highest level of sequence polymorphism between *Pyricularia oryzae* and the other *Pyricularia* species, with a greater number of variable sites observed across the alignment. Specificity was evaluated by dPCR using 19 strains of *P. oryzae* from various hosts (Table S1). The designed primers and probe enabled successful amplification of all *P. oryzae* strains tested. A total of 36 non-target strains, belonging to the genera *Curvularia, Pseudopyricularia, Proxipyricularia* and to other *Pyricularia* species than *P. oryzae*, were also analyzed (Table S1). Among these, only two strains were amplified by dPCR with the developed oligonucleotides. These strains belong to the species *P. ctenantheicola* (GR0001 and GR0002). However, the fluorescence intensity observed for *P. ctenantheicola* strains was lower and close to the detection threshold (Figure 2).

**Figure 2.**
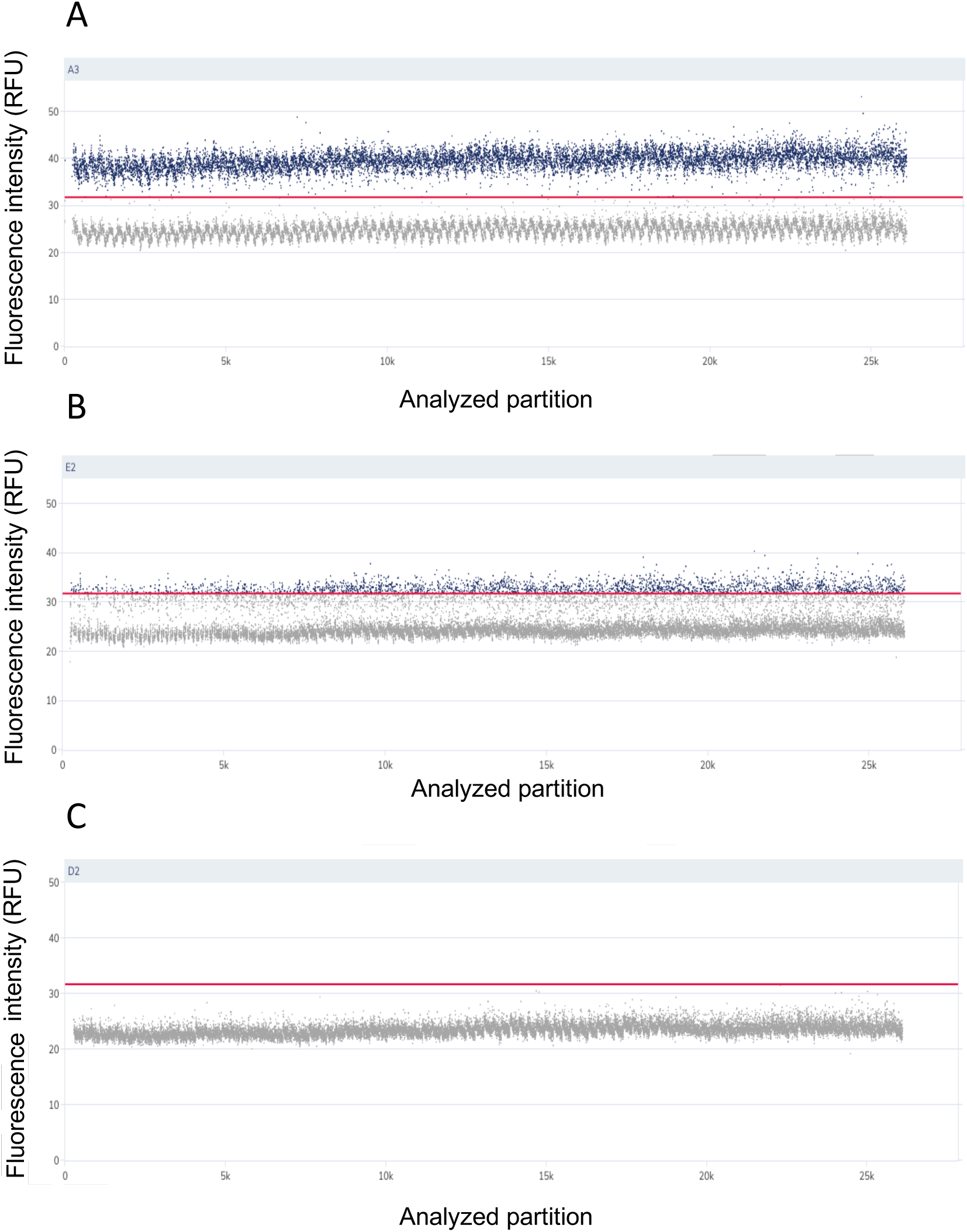
Raw dPCR results from amplification of *Pyricularia* spp. pure DNA. Example of results obtained for a *Pyricularia oryzae* strain (A). Example of results obtained for a *Pyricularia ctenantheicola* strain (B). Example of results obtained for the 33 strains belonging to different species of *Pyricularia* spp. (other than *P. oryzae*), *Pseudopyricularia* spp., *Curvularia* spp., and *Helminthosporium* spp. (C). Each point represents one dPCR reaction mix partition. Gray points represent partitions determined as negative (no PCR amplification), and blue points represent partitions determined as positive (PCR amplification), according to their fluorescence intensity (RFU). The red line, indicating the threshold separating positive and negative partitions, is placed **a**fter comparison between positive and negative controls.

### 3.2. *P. oryzae* is detected early after inoculation and DNA concentration stabilizes over time

The infection dynamic of *P. oryzae* was evaluated by quantifying fungal DNA over time, after inoculation of perennial ryegrass. Tukey’s post-hoc test revealed a significant increase of *P. oryzae* abundance between day 0 and day 5, from 4 x 10^3^ to 1 x 10^7^ copies of DNA.g of sample^-1^ (*p* < 0.001) (Figure 3A). During this period, *ie* three days after inoculation, the first symptoms appeared. Between day 5 and day 10, *P. oryzae* abundances detected in the samples were not significantly different (*p* > 0.05; average 1.3 x 10^7^). Indeed, as shown in Figure 3A, the quantification remained stable over the last three sampling points, suggesting that *P. oryzae* DNA accumulation reached a plateau five days after inoculation.

**Figure 3.**
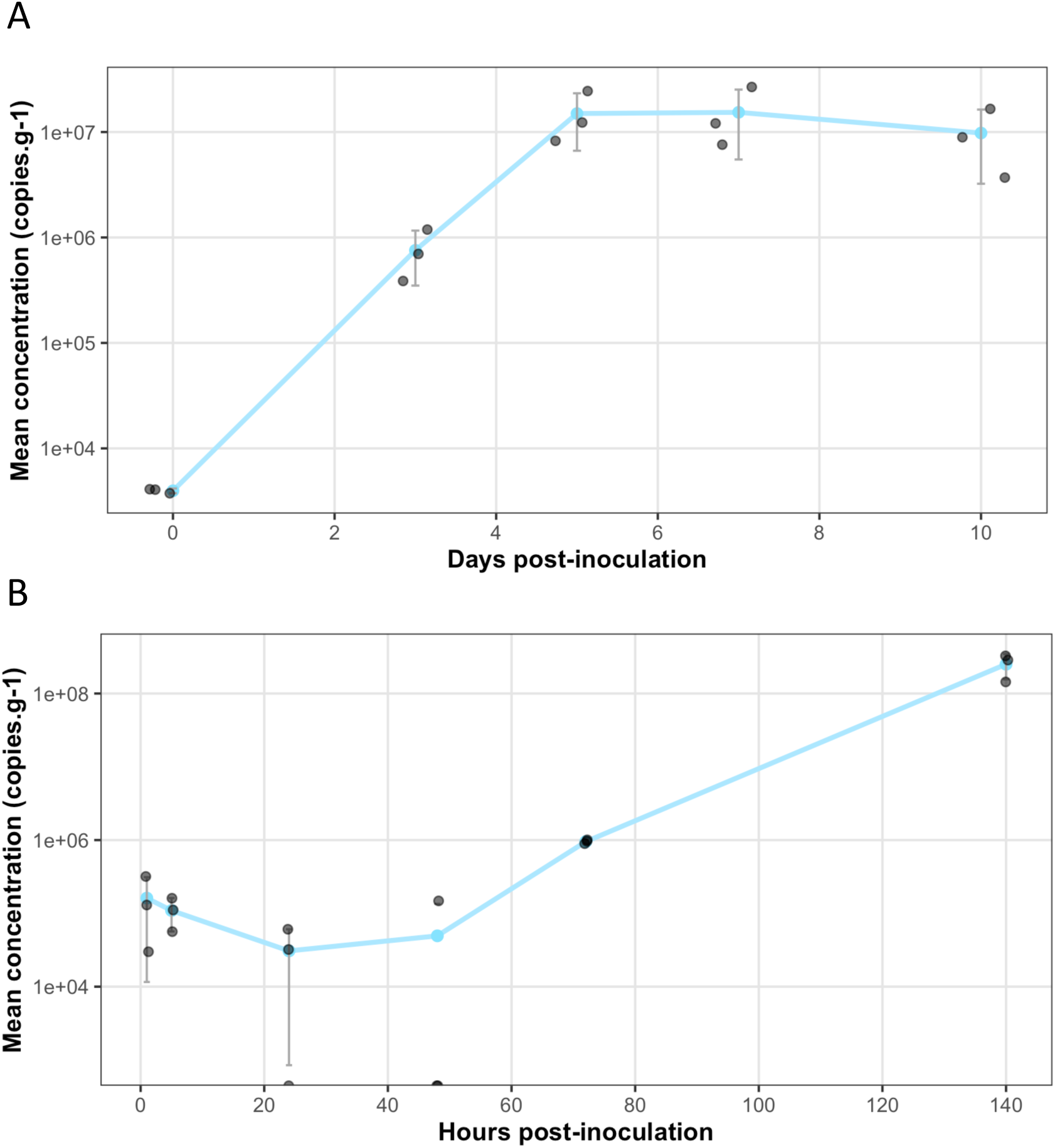
Temporal dynamics of *P. oryzae* DNA accumulation in inoculated grass samples. (A) Experiment 1: 0-10 days. (B) Experiment 2: 1-140h. Mean *P. oryzae* DNA quantities (copies.g^-1^, ± standard deviation, n = 3) are plotted against time after inoculation.

To refine this dynamic and determine the earliest detection time, a second experiment was performed with earlier and shorter sampling intervals (hours post inoculation instead of days). Overall, similarly to the first experiment, *P. oryzae* DNA levels increased from approximately 10^5^ to 10^7^–10^8^ copies of DNA.g^-1^ over the course of the experiment, with the same quantity of DNA measured for both experiments at day 3 and after day 5. However, during the first 24 h post-inoculation, DNA levels showed a slight decrease before progressively increasing at later time points.

An overall significant effect of time was observed (one-way ANOVA. F-value = 5.81; *p* = 0.006). Tukey’s post-hoc test revealed that DNA quantity was higher at 140h (ca. 6 days) post-inoculation than at 24h and 48 h post-inoculation. However, no significant differences were observed among the other sampling time points (p > 0.05). *Pyricularia oryzae* multiplication within the plant showed a progressive increase after 24h post-inoculation, and the increase became significant at 140 h post-inoculation (Figure 3B).

### 3.3. *Pyricularia oryzae* is detected days before symptom appearance and remains detectable long after the disease outbreak

*P. oryzae* was not detected by the classical method of microscopic observation of sporulation from 05/12/2022 to 26/06/2023 included. Starting on 03/07/2023, the number of plants producing spores after incubation increased rapidly, showing that an epidemic was in progress. With the dPCR method, *P. oryzae* was detected in aerial parts of turf at almost all sampling dates between 05/12/2022 and 12/06/2023. A slight increase of *P. oryzae* DNA quantity, between 01/05/2023 and 12/06/2023, is probably indicative of the beginning of the gray leaf spot epidemic (Figure 4). The epidemic development was also confirmed by relatively high and increasing quantities of *P. oryzae* DNA in mowing debris from the same stadium: 1.1, 1.7 and 5.6 x 10^5^ DNA-copies.g^-1^ on July 7, 11 and 18, respectively (data not shown). So, the dPCR method allowed (*i*) to detect the fungus in the winter and spring seasons, when disease symptoms are not observed, and (*ii*) to detect the premises of the epidemic three weeks before the detection by the classical method.

**Figure 4.**
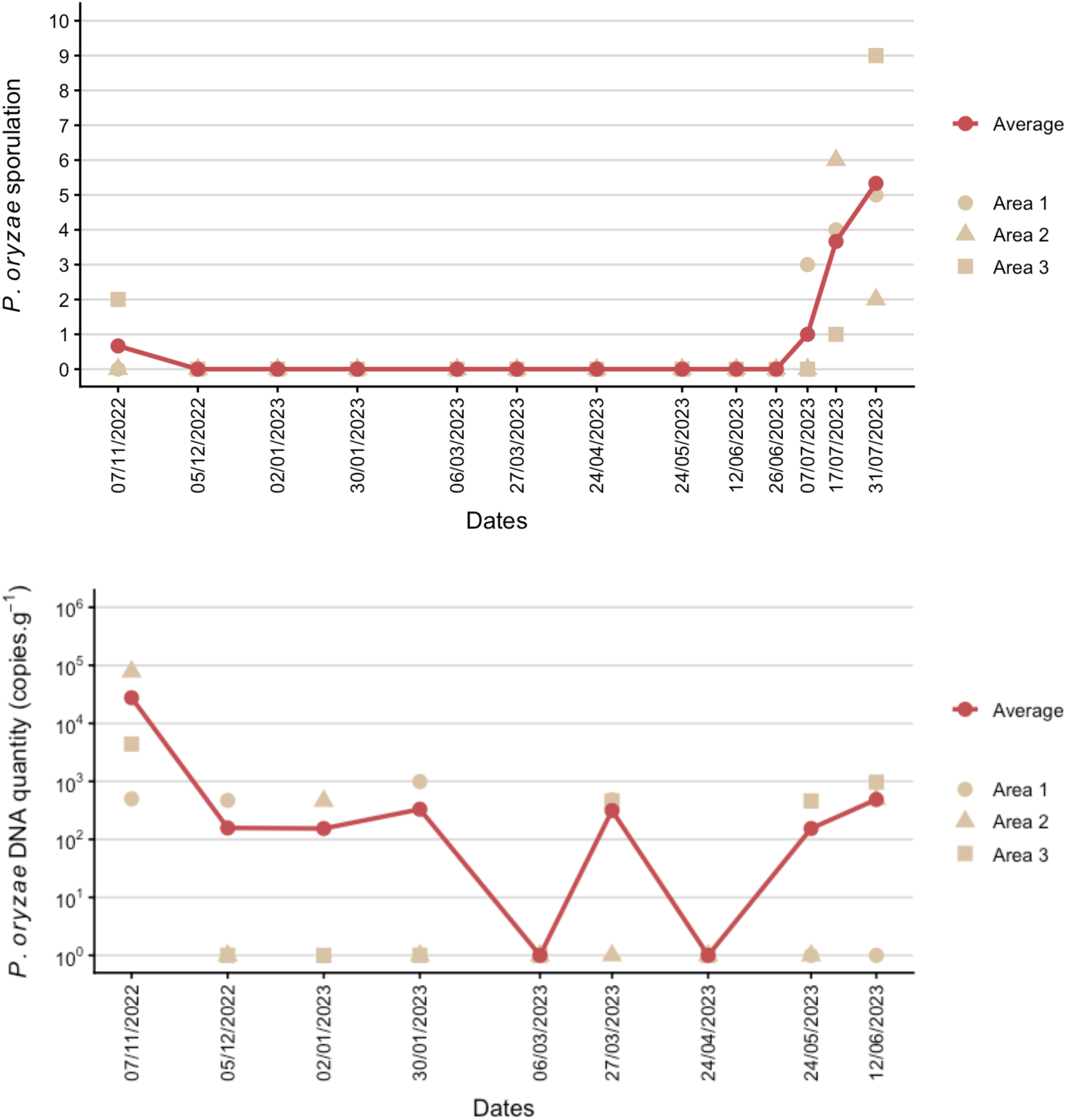
Long term survey of *P. oryzae* in aerial parts of turf on a stadium. (A) Detection by microscopic observation. Sporulation was measured by counting the number of sporulating lesions on the samples observed. (B) Detection by digital PCR.

### 3.4. The method detects *P. oryzae* at low proportions of infected plants

Infected perennial ryegrass was mixed with different quantities of healthy perennial ryegrass to characterize the limit of detection (defined as the smallest dilution at which *P. oryzae* was detected in all three replicates)*. P. oryzae* was not detected in the negative control (0% of infected grass) and was quantified at 1.47 x 10^7^ DNA.g of samples^-1^ in the positive control (100% of infected grass). The fungus was detected in all three replicates at 0.1% and 0.05% infected sample. At 0.025%, only 2 out of 3 replicates were positive (Table S2). The detection limit of the method was therefore defined as 0.05% of infected plants corresponding to a mean concentration of 1.3 x 10^4^ copies of DNA.g^-1^ of samples.

### 3.5. *Pyricularia oryzae* is detected in the soil, in the roots and in the aerial parts of turf in stadium

The presence of *P. oryzae* in different matrices was evaluated by quantifying fungal DNA in the soil, roots and aerial parts of plants collected from 15 stadiums during spring. *P. oryzae* was detected in at least one of the three matrices in 13 out of the 15 stadiums analyzed, indicating a high prevalence across the surveyed stadiums. The results for each matrix are presented in Figure 5 (Table S3). Among the 15 stadiums, *P. oryzae* was frequently detected: in 9 soil samples, 10 root samples and 5 aerial parts of turf samples. Roots therefore exhibited the highest prevalence among the three matrices.

**Figure 5.**
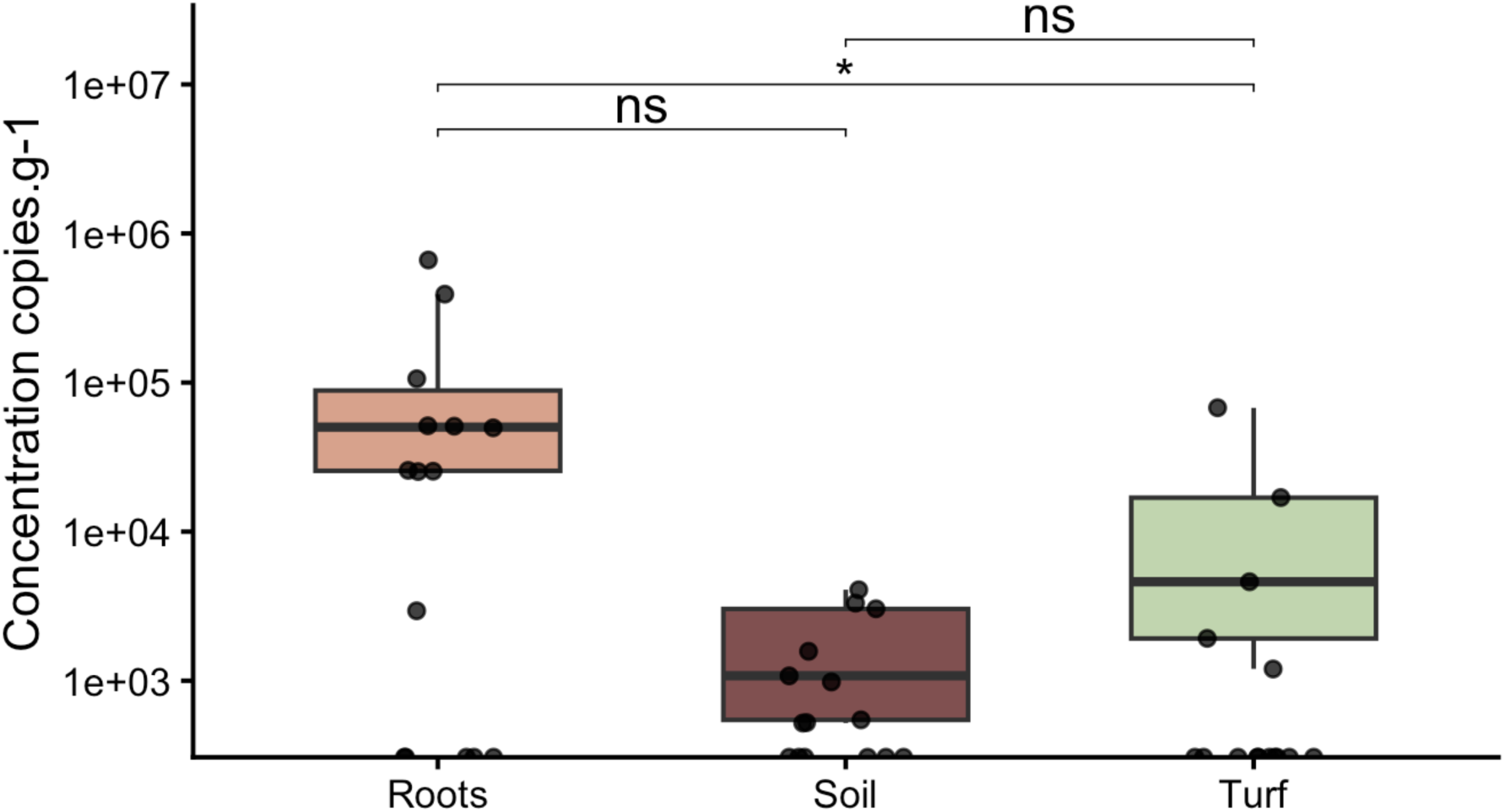
Quantity of *P. oryzae* DNA in soil, roots sand aerial parts of turf. Boxplots represent median and interquartile range; points represent individual stadiums. Statistical correlation between matrices was assessed using Pairwise Spearman rank correlations (n=15). Significant correlation are indicated by an asterisk (p < 0.05); ns: non significative.

A Friedman test revealed significant differences in *P. oryzae* abundance among the three matrices (*p* = 0.0069). Root samples showed the highest mean for *P. oryzae* DNA abundance (even when negative samples were included and set to zero; Figure 5). Pairwise Wilcoxon signed-rank tests with Holm correction showed that *P. oryzae* DNA abundance was significantly higher in roots compared with soil (*p* = 0.018) and aerial parts (*p* = 0.018), whereas no significant difference was observed between soil and aerial parts samples (*p* = 0.919). Pairwise Spearman rank correlations were then performed to investigate whether the concentration of *P. oryzae* was correlated between matrices. No significant correlation was observed between soil and roots (Spearman’s ρ = −0.045, *p* = 0.873), or between soil and aerial parts (Spearman’s ρ = 0.188, p = 0.503). However, a significant correlation was observed between roots and aerial parts (Spearman’s ρ = 0.576, *p* = 0.025), suggesting that *P. oryzae* abundance is linked between the two matrices.

### 3.6. *Pyricularia oryzae* is widely distributed across the playing field of a stadium

In the roots compartment, *P. oryzae* was detected in 27 out of 32 (94.4%) areas sampled in the stadium analyzed. In these infected zones, concentrations of *P. oryzae* DNA ranged from 4.91 x 10^2^ to 8.18 x 10^4^ copies of DNA.g^-1^, with 9 zones exceeding 10^4^ copies.g^-1^. The highest infection levels were concentrated in the center of the field (Figure 6). In the aerial compartment, only five zones (15.6%) were tested positive for *P. oryzae*, with lower concentrations than for roots, ranging from 6.5 x 10^2^ to 2.07 x 10^3^ copies of DNA.g^-1^. When both compartments are considered together, 29 out of 32 zones were affected by *P. oryzae*, corresponding to 90.6% of the field.

**Figure 6.**
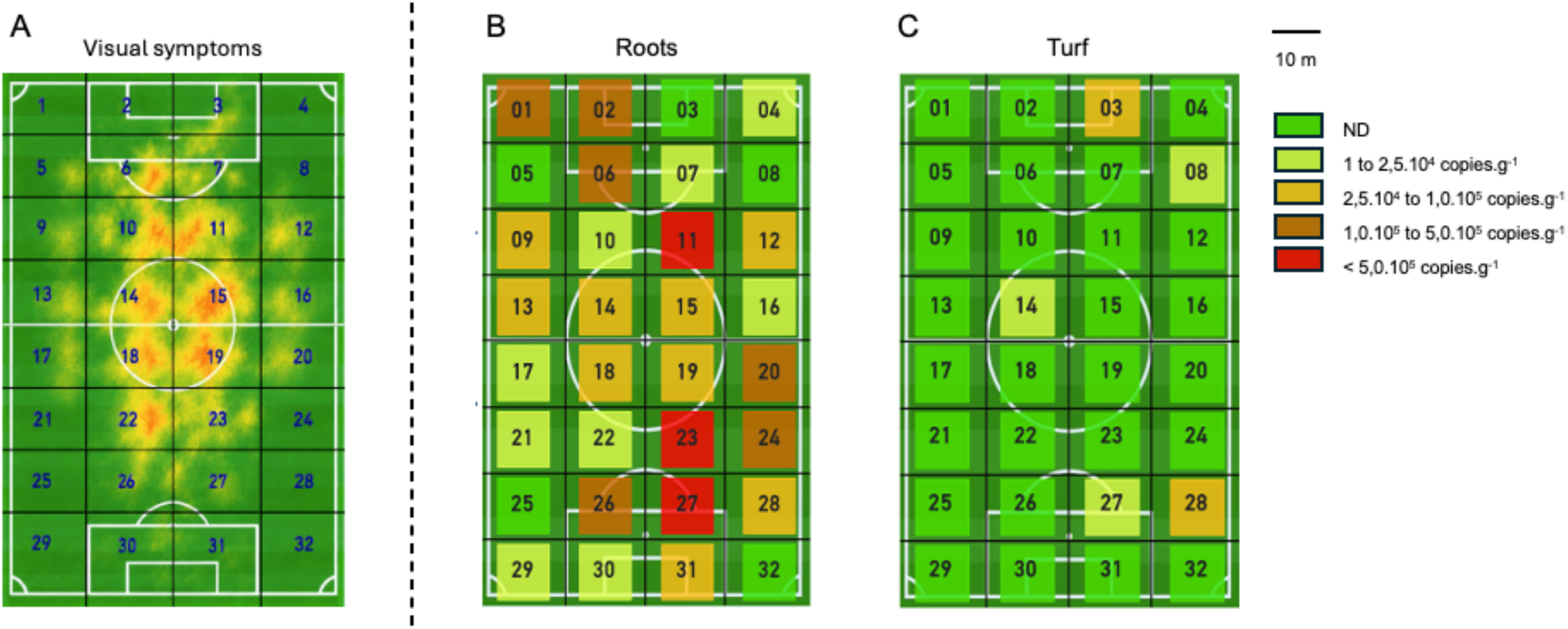
Cartography of *P. oryzae* detection on the 32 zones delimited on the stadium. (A) Schematic representation of the visual symptoms during the epidemic (29/08/2024) (B) Schematic representation of the levels of DNA quantification of *P. oryzae* in the root compartment sampled on the 20/11/2024 (C) Schematic representation of the levels of DNA quantification of *P. oryzae* in the turf compartment sampled on the 20/11/2024. The schematic representation of the visual symptoms was generated with the assistance of ChatGPT (OpenAI), based on field observations and photographs provided by the authors.

## Discussion

Accurate and early diagnostic tools are essential for the development of effective disease management strategies. In turfgrass, gray leaf spot caused by *P. oryzae,* represents an increasing concern, particularly in intensively managed turf such as sports fields, where favorable environmental conditions can lead to rapid disease development and severe losses in turf quality (Uddin et al. 2003a). Indeed, professional sports stadiums represent highly managed and dynamic ecosystems where disease, the interaction between the pathogen and the host plant, is favored by intensive cultural practices, and specific environmental conditions created within the stadium. Management of gray leaf spot remains challenging because curative control is often limited when extensive colonization and symptom development have occurred (Kim et al. 2003). Stadiums require a surveillance strategy that goes beyond disease detection to monitor the spatio-temporal circulation of the pathogen, enabling both early disease detection and new epidemiological insights for an integrated disease management. Preventive and early intervention strategies are considered the most effective approaches for reducing disease impact (Kim et al. 2003). Taken together, our results indicate that multi-matrix dPCR extends gray leaf spot diagnosis from symptomatic confirmation to spatio-temporal surveillance at stadium scale. The assay enabled pre-symptomatic detection of *P. oryzae* and quantitative monitoring across soil, roots and aerial tissues, including during asymptomatic periods and after an outbreak.

While molecular tools have considerably improved the detection of *P. oryzae*, they have mostly been developed as diagnostic methods applied to symptomatic plant tissues. In this context, the development of a multi-matrix, specific and sensitive dPCR approach represents an important step toward improving disease monitoring and optimizing treatment timing. Early molecular detection is not unique to dPCR as previous PCR and LAMP-based approaches have demonstrated the potential for rapid and early detection of *P. oryzae* in turfgrass (Harmon et al. 2003, Villari et al. 2017). The main contribution of the present study therefore lies not only in early detection, but in the integration of quantitative measurements across several stadium compartments. In contrast with methods primarily designed for laboratory confirmation or single-matrix monitoring, the strategy provides complementary information on underlying pathogen presence, temporal changes and stadium heterogeneity.

The developed primers and probe showed broad inclusivity, as all nineteen *P. oryzae* isolates included in the validation panel were successfully detected. Thirty-four of the thirty-six non-target isolates were negative, indicating a high level of analytical discrimination. The two exceptions were strains of *P. ctenantheicola*, but the amplification of *P. oryzae* and *P. ctenantheicola* can be distinguished based on fluorescence intensity pattern of the positive clusters, allowing reliable discrimination between the two species. Given that *P. ctenantheicola* has not been reported as a turfgrass pathogen, and its known hosts are taxonomically distinct from commonly used turfgrasses, this limited cross-reactivity is unlikely to compromise the assay’s practical application.

The first experiment in controlled conditions indicated that *P. oryzae* DNA accumulation mainly occurred during the first five days following inoculation. To further characterize this early phase of infection, a second experiment was conducted on the firsts hours post-inoculation. Interestingly, although *P. oryzae* was detected immediately after inoculation, DNA quantity decreased during the first 24h. This pattern suggests that a substantial fraction of the initial signal may correspond to residual inoculum deposited on the leaf surface, rather than to fungal multiplication within plant tissues. The relatively high variability between samples supports this hypothesis. The subsequent increase in fungal DNA after 24h is more consistent with the onset of a progressive multiplication of *P. oryzae* in plant tissue. Fungus could be detected in the hours following inoculation, long before symptoms became visible (3 days), supporting pre-symptomatic detection. However, the onset of active colonization could not be determined more precisely because DNA quantities detected were not significantly different between 24 and 48h. Early detection was confirmed by experiments in stadium real conditions. The beginning of the epidemic was detected three weeks before the detection by the classical method. The dPCR detection performed better in terms of earliness compared to classical methods, and this is likely due to method sensitivity.

Indeed, for practical implementation at the stadium scale, a simple sampling strategy could consist of collecting mown grass representative of the entire field. In such a composite sample, infected tissues may represent only a very small proportion of the total material, making the sensitivity of the detection method particularly important. To assess sensitivity, decreasing proportions of infected grass mixed with healthy grass were tested. *P. oryzae* was detected in all replicates at the proportion 0.05% infected tissue tested. This demonstrates that the method remains highly sensitive even when infected tissue represents only a very small fraction of the total sample. Although these experiments were conducted under controlled conditions and do not fully reproduce the heterogeneity and scale of a stadium, the results provide encouraging evidence for the detection of low proportions of infected tissue within composite samples. This sensitivity therefore supports the potential use of pooled grass clippings as a simple and practical sampling strategy for large-scale monitoring and early detection of *P. oryzae* in stadium turfgrass.

It is also noteworthy that low levels of *P. oryzae* DNA were detected by dPCR in samples collected from November to June, when environmental conditions are less favorable for disease development. This persistent detection indicates that *P. oryzae* DNA remains detectable with dPCR in the stadium throughout the unfavorable season. Although DNA detection alone does not demonstrate pathogen viability, these observations raise the possibility that *P. oryzae* may persist between disease-favorable periods and contribute to renewed disease development when environmental conditions become favorable again.

Although *P. oryzae* is primarily associated with foliar disease, other plant compartments may contribute to its persistence or provide useful matrices for surveillance. To investigate this possibility and identify the most relevant matrix for diagnostic purposes, soil, roots and aerial tissues were analyzed across 15 stadiums sampled in spring, before environmental conditions became favorable for disease development. *P. oryzae* was detected in all three matrices despite the absence of visible symptoms. Roots showed the highest prevalence and significantly higher fungal abundance than soil or aerial parts. Furthermore, fungal abundance in roots and aerial tissues was positively correlated, whereas no significant correlation involved the soil compartment. These results identify roots as the most informative matrix in this dataset for detecting *P. oryzae* during asymptomatic periods. Establishing roots as a biologically active reservoir remains to be demonstrated. Quantitative comparisons among matrices should also be interpreted cautiously because sample masses, pretreatments, and extraction procedures differed, and DNA recovery may be influenced by extraction efficiency, matrix inhibition, and the spatial heterogeneity of fungal material. Moreover, only 15 stadiums were included in this study, and several factors were uncontrolled across sites, including disease management practices, turfgrass maintenance, and the cultivars used (which may differ in their susceptibility to gray leaf spot). These factors could influence *P. oryzae* development and consequently the amount of fungal DNA detected in the different matrices. Nevertheless, in spring, root sampling could therefore complement aerial-tissue monitoring when foliar DNA quantities are low or visible symptoms are absent and help anticipate periods of increased disease risk.

Furthermore, when investigating the spatial distribution of *P. oryzae* within a stadium following a gray leaf spot outbreak, *P. oryzae* was widely detected in the root, with 27 positive samples out of the 32 zones tested, whereas only 5 of the 32 zones were positive with aerial tissues. The selected stadium had developed visible symptoms during the summer and was subsequently subjected to disease management practices, including treatment and overseeding. Sampling was performed several months later, after visible symptoms had disappeared. Interestingly, comparison with the distribution of symptoms recorded during the previous outbreak showed an apparent spatial correspondence between previously symptomatic areas and zones in which *P. oryzae* DNA was subsequently detected in roots (Figure 6). This observation raises the possibility that the pathogen may persist in the root compartment in areas previously affected by the disease, which could potentially contribute to disease recurrence in these areas when environmental conditions become favorable again. At the same time, the widespread detection of *P. oryzae* across the stadium indicates that its presence was not restricted to previously symptomatic areas. This broad distribution may reflect pathogen dispersal within the stadium, potentially facilitated by turf management practices, although the mechanisms underlying this distribution remain to be determined. However, as this spatial pattern was observed in a single stadium, further investigations are needed to determine whether it can be generalized to other stadiums. Overall, the persistence of *P. oryzae* DNA in asymptomatic turf several months after the outbreak highlights the potential of root sampling for detecting the pathogen beyond periods of visible disease.

In conclusion, combining molecular quantification with epidemiological and management-related factors could contribute to the development of more reliable tools for anticipating gray leaf spot outbreaks in stadium turfgrass. Repeated sampling of aerial tissues, including pooled mowing debris, could reveal changes in *P. oryzae* DNA before extensive symptoms develop, while spatial sampling could identify areas requiring closer inspection. Together, these measurements could help guide the timing and location of preventive measures, including irrigation adjustments, turf management, and biocontrol applications. To this aim, the relationship between *P. oryzae* DNA levels and subsequent disease development under stadium conditions still requires further investigation to determine whether specific DNA concentrations could be used as indicators of increased disease risk. Moreover, screening incoming plant material, as seed lots or turf rolls could therefore complement stadium monitoring as a preventive approach to reduce the risk of introducing *P. oryzae* and, consequently, the risk of gray leaf spot development. For example, *P. oryzae* DNA has already been detected in seed samples using the method developed in this study (data not shown), although this detection does not demonstrate pathogen viability or seed transmission. This finding is consistent with previous reports of seedborne detection of *P. oryzae* in rice seeds (Manandhar et al., 1998; Thierry et al., 2020b). *P. oryzae* quantification can also provide a complementary endpoint for trials of biocontrol agents and other integrated management practices. Measurements before intervention and at predefined intervals afterwards would help describe the dynamics associated with treatment. Where a microbial biocontrol agent is studied, its establishment could be measured separately with an agent-specific assay. Such studies are needed to determine whether combining molecular measurements with field observations improves the timing and assessment of integrated gray leaf spot management.

## Notes

### Competing Interest Statement

The authors have declared no competing interest.

## References

1. Baudin, M., Le Naour-Vernet, M., Gladieux, P., Tharreau, D., Lebrun, M.-H., Lambou, K., Leys, M., Fournier, E., Césari, S., & Kroj, T. (2024). *Pyricularia oryzae*: Lab star and field scourge. Molecular Plant Pathology, 25(4), e13449. 10.1111/mpp.13449

2. Bonos, S. A., Clarke, B. B., & Murphy, J. A. (2006). Integrated control of gray leaf spot on perennial ryegrass. Rutgers NJAES Cooperative Extension, FS1048.

3. Couch, B. C., Fudal, I., Lebrun, M.-H., Tharreau, D., Valent, B., van Kim, P., Nottéghem, J.-L., & Kohn, L. M. (2005). Origins of host-specific populations of the blast pathogen *Magnaporthe oryzae* in crop domestication with subsequent expansion of pandemic clones on rice and weeds of rice. Genetics, 170(2), 613–630. 10.1534/genetics.105.041780

4. Farman, M. L. (2002). *Pyricularia grisea* isolates causing gray leaf spot on perennial ryegrass (*Lolium perenne*) in the United States: Relationship to *P. grisea* isolates from other host plants. Phytopathology, 92(3), 245–254. 10.1094/PHYTO.2002.92.3.245

5. Gladieux, P., Ravel, S., Rieux, A., Cros-Arteil, S., Adreit, H., Milazzo, J., Thierry, M., Fournier, E., Terauchi, R., & Tharreau, D. (2018). Coexistence of multiple endemic and pandemic lineages of the rice blast pathogen. mBio, 9(2), e01806–17. 10.1128/mBio.01806-17

6. Groben, G., Clarke, B. B., Murphy, J., Koch, P., Crouch, J. A., Lee, S., & Zhang, N. (2020). Real-time PCR detection of *Clarireedia* spp., the causal agents of dollar spot in turfgrasses. Plant Disease, 104(12), 3118–3123. 10.1094/PDIS-04-20-0726-RE

7. Harmon, P. F., Dunkle, L. D., & Latin, R. (2003). A rapid PCR-based method for the detection of *Magnaporthe oryzae* from infected perennial ryegrass. Plant Disease, 87(9), 1072–1076. 10.1094/PDIS.2003.87.9.1072

8. Huang, W., Liu, X., Zhou, X., Wang, X., Liu, X., & Liu, H. (2020). Calcium signaling is suppressed in *Magnaporthe oryzae* conidia by *Bacillus cereus* HS24. Phytopathology, 110(2), 309–316. 10.1094/PHYTO-08-18-0311-R

9. Kim, Y. S., Dixon, E. W., Vincelli, P., & Farman, M. L. (2003). Field resistance to strobilurin (QoI) fungicides in *Pyricularia grisea* caused by mutations in the mitochondrial cytochrome b gene. Phytopathology, 93(7), 891–900. 10.1094/PHYTO.2003.93.7.891

10. Li, L., Zhang, S. Y., & Zhang, C. Q. (2019). Establishment of a rapid detection method for rice blast fungus based on one-step loop-mediated isothermal amplification (LAMP). Plant Disease, 103(8), 1967–1973. 10.1094/PDIS-11-18-1964-RE

11. Manandhar, H. K., Jorgensen, H. L., Smedegaard-Petersen, V., & Mathur, S. B. (1998). Seedborne infection of rice by Pyricularia oryzae and its transmission to seedlings. Plant Disease, 82(10), 1093–1099. 10.1094/PDIS.1998.82.10.1093

12. Milazzo, J., Pordel, A., Ravel, S., & Tharreau, D. (2019). First scientific report of *Pyricularia oryzae* causing gray leaf spot disease on perennial ryegrass (*Lolium perenne*) in France. Plant Disease, 103(5), 1024. 10.1094/PDIS-09-18-1545-PDN

13. Pordel, A., Ravel, S., Charriat, F., Gladieux, P., Cros-Arteil, S., Milazzo, J., Adreit, H., Javan-Nikkhah, M., Mirzadi-Gohari, A., Moumeni, A., & Tharreau, D. (2021). Tracing the origin and evolutionary history of *Pyricularia oryzae* infecting maize and barnyard grass. Phytopathology, 111(1), 128–136. 10.1094/PHYTO-09-20-0423-R

14. Tanaka, A., Tamba, H., Tosa, Y., Mayama, S., & Tani, T. (2000). Grey leaf spot occurred on perennial ryegrass (*Lolium perenne*) on golf courses (2). Journal of Japanese Society of Turf Science, 29, 38–39. [Abstract in Japanese].

15. Thierry, M., Gladieux, P., Fournier, E., Tharreau, D., & Ioos, R. (2020a). A genomic approach to develop a new qPCR test enabling detection of the *Pyricularia oryzae* lineage causing wheat blast. Plant Disease, 104(1), 60–70. 10.1094/PDIS-04-19-0685-RE

16. Thierry, M., Chatet, A., Fournier, E., Tharreau, D., & Ioos, R. (2020b). A PCR, qPCR, and LAMP toolkit for the detection of the wheat blast pathogen in seeds. Plants, 9(2), 277. 10.3390/plants9020277

17. Tosa, Y., Uddin, W., Viji, G., Kang, S., & Mayama, S. (2007). Comparative genetic analysis of *Magnaporthe oryzae* isolates causing gray leaf spot of perennial ryegrass turf in the United States and Japan. Plant Disease, 91(5), 517–524. 10.1094/PDIS-91-5-0517

18. Tredway, L. P., Stevenson, K. L., & Burpee, L. L. (2005). Genetic structure of *Magnaporthe grisea* populations associated with St. Augustinegrass and tall fescue in Georgia. Phytopathology, 95(5), 463–471. 10.1094/PHYTO-95-0463

19. Uddin, W., Viji, G., & Vincelli, P. (2003a). Gray leaf spot (blast) of perennial ryegrass turf: An emerging problem for the turfgrass industry. Plant Disease, 87(8), 880–889. 10.1094/PDIS.2003.87.8.880

20. Uddin, W., Viji, G., Schumann, G. L., & Boyd, S. H. (2003b). Detection of *Pyricularia grisea* causing gray leaf spot of perennial ryegrass turf by a rapid immuno-recognition assay. Plant Disease, 87(7), 772–778. 10.1094/PDIS.2003.87.7.772

21. Uddin, W., Serlemitsos, K., & Viji, G. (2003c). A temperature and leaf wetness duration-based model for prediction of gray leaf spot of perennial ryegrass turf. Phytopathology, 93(3), 336–343. 10.1094/PHYTO.2003.93.3.336

22. Viji, G., Wu, B., Kang, S., Uddin, W., & Huff, D. R. (2001). *Pyricularia grisea* causing gray leaf spot of perennial ryegrass turf: Population structure and host specificity. Plant Disease, 85(8), 817–826. 10.1094/PDIS.2001.85.8.817

23. Villari, C., Mahaffee, W. F., Mitchell, T. K., Pedley, K. F., Pieck, M. L., & Peduto Hand, F. (2017). Early detection of airborne inoculum of *Magnaporthe oryzae* in turfgrass fields using a quantitative LAMP assay. Plant Disease, 101(1), 170–177. 10.1094/PDIS-06-16-0834-RE

24. Vincelli, P. (2000). Fungicidal control of gray leaf spot: Disease need not destroy perennial ryegrass. Golf Course Management, 68, 68–74.

25. Vincelli, P., & Dixon, E. (2002). Resistance to QoI (strobilurin-like) fungicides in isolates of *Pyricularia grisea* from perennial ryegrass. Plant Disease, 86(3), 235–240. 10.1094/PDIS.2002.86.3.235

26. Vines, P. L., Daddio, R. M., Luo, J., Wang, R., Murphy, J. A., Zhang, N., Clarke, B. B., Meyer, W. A., & Bonos, S. A. (2022). *Pyricularia oryzae* incites gray leaf spot disease on hard fescue (*Festuca brevipila*). International Turfgrass Society Research Journal, 14, 997–1002. 10.1002/its2.17

27. Williams, D. W., Burrus, P. B., & Vincelli, P. (2001). Severity of gray leaf spot in perennial ryegrass as influenced by mowing height and nitrogen level. Crop Science, 41, 1207–1211. 10.2135/cropsci2001.4141207x

