## Supplementary tables 1 2 3 for "Towards operational epidemiology of *Pyricularia oryzae* in sports turf: contribution of a multi-matrix molecular monitoring strategy at stadium scale"

### Supplementary data

**Table S1. List of microorganisms used for the design and specificity tests of primers and probes targeting *Pyricularia oryzae***

| Species | Strain number | Host | Sample localisation | Design and in silico specificity | In vitro specificity and inclusivity | Detection dPCR |
|---|---|---|---|---|---|---|
| *Pyricularia oryzae* | FR1080 | *Lolium perenne* | France, Montpellier | | x | + |
| | FR1081 | *Lolium perenne* | France, Montpellier | | x | + |
| | FR1082 | *Lolium perenne* | France, Montpellier | | x | + |
| | FR1083 | *Lolium perenne* | France, Montpellier | | x | + |
| | FR1084 | *Lolium perenne* | France | | x | + |
| | FR1086 | *Lolium perenne* | France | | x | + |
| | FR1090 | *Lolium perenne* | France | | x | + |
| | DE0001 | *Lolium perenne* | Germany | x | | |
| | FR1067 | *Lolium perenne* | France, Montpellier | x | | |
| | FR1142 | *Lolium perenne* | France | x | | |
| | PgKY4OV2.1 | *Lolium perenne* | USA, Kentucky, Lexington | x | | |
| | AT0011 | *Lolium perenne* | Austria | x | | |
| | AG0062 | *Lolium sp.* | Argentina, Corrientes | x | x | + |
| | AG0063 | *Lolium sp.* | Argentina, Corrientes | | x | + |
| | AG0064 | *Lolium sp.* | Argentina, Corrientes | | x | + |
| | CR0026 | *Lolium hybridum* | Republic of Korea, Gyeonggi, Suweon | | x | + |
| | AG0102 | *Triticum aestivum* | Argentina, Concepción del Uruguay | x | | |
| | BTGP-1(b) | *Triticum aestivum* | Bangladesh, Meherpur, Gangni | x | | |
| | B71 | *Triticum aestivum* | Bolivia, Okinawa Uno | x | | |
| | Br80 | *Triticum aestivum* | Brazil | x | | |

| | | | | | | |
|---|---|---|---|---|---|---|
| | PY0925 | *Triticum aestivum* | Brazil, Minas Gerais, Perdizes | x | | |
| | PY6017 | *Triticum aestivum* | Brazil, Coromandel | x | | |
| | ZMW18_06 | *Triticum aestivum* | Zambia | x | | |
| | BR0032 | *Triticum sp.* | Brazil | | x | + |
| | IN0017 | *Oryza sativa* | India | x | x | + |
| | BF0072 | *Oryza sativa* | Burkina Faso, Pankatorio | x | | |
| | CH0533 | *Oryza sativa* | China,Yunnan, Baoshan | x | | |
| | CH0999 | *Oryza sativa* | China,Yunnan,Yule | x | | |
| | CH1897 | *Oryza sativa* | China,Yunnan, Xiaoshuijng | x | | |
| | CL3-6-7 | *Oryza sativa* | Colombia, Santa Rosa | x | | |
| | IT0010 | *Oryza sativa* | Italy, Veneria | x | | |
| | LA0003 | *Oryza sativa* | Laos, Luang prabang, Silalek | x | | |
| | JP0038 | *Eragrostis curvula* | Japan | x | x | + |
| | JP0027 | *Eragrostis curvula* | Japan | x | | |
| | G17 | *Eragrostis curvula* | Japan | x | | |
| | JP0032 | *Eragrostis lehmannia* | Japan | x | | |
| | BR0070 | *Eragrostis sp.* | Brazil, Mato Grosso | x | x | + |
| | BR0072 | *Eragrostis sp.* | Brazil, Mato Grosso | x | | |
| | PH0078 | *Echinochloa sp.* | Philippines | | x | + |
| | IR0083 | *Echinochloa sp.* | Iran, Golestan, Agh Ghala | x | | |
| | IR0088 | *Echinochloa sp.* | Iran, Golestan, Agh Ghala | x | | |
| | IR0102 | *Echinochloa sp.* | Iran, Golestan, Azadshahr | x | | |
| | IR0109 | *Setaria galica* | Iran, Golestan, Gharakhil | x | | |
| | GFSI1-7-2 | *Setaria italica* | Japan,Gifu | x | | |
| | IN0007 | *Setaria italica* | India | x | x | + |
| | SV9610 | *Setaria viridis* | China, Zhejiang | x | | |
| | SV9623 | *Setaria viridis* | China, Zhejiang | x | | |

| | | | | | | |
|---|---|---|---|---|---|---|
| | US0064 | *Setaria sp.* | USA, Maryland | x | | |
| | OG0002 | *Eleusine coracana* | Uganda | x | | |
| | JP0015 | *Eleusine coracana* | Japan | x | | |
| | EI9411 | *Eleusine indica* | China, Fujian | x | | |
| | EI9604 | *Eleusine indica* | China, Zhejiang | x | | |
| | PH0075 | *Brachiara mutica* | Philippines | x | x | + |
| | PH0035 | *Brachiaria mutica* | Philippines, Los Banos | x | | |
| | PH0045 | *Brachiaria mutica* | Philippines | x | | |
| | PH0046 | *Brachiaria distachya* | Philippines | x | | |
| | PR0083 | *Stenotaphrum secundatum* | Portugal, Elvas | x | | |
| | PR0104 | *Stenotaphrum secundatum* | Portugal, Samora Correia | x | | |
| | AG0065 | *Stenotaphrum sp.* | Argentina, Concepción del Uruguay | x | | |
| | GN0001 | *Zea mays* | Gabo | x | x | + |
| | IR0095 | *Zea mays* | Iran, Golestan, Agh Ghala | | | |
| **Other Pyricularia species** | | | | | | |
| *Pyricularia ctenantheicola* | GR0001 | *Ctenanthe oppenheimiana* | Greece, Volos | x | x | + |
| | GR0002 | *Ctenanthe oppenheimiana* | Greece, Volos | | x | + |
| *Pyricularia grisea* | BR0029 | *Digitaria sanguinalis* | Brazil, Goias, Goiana | x | x | - |
| | JP0034 | *Digitaria smutsii* | Japan | | x | - |
| | PH0055 | *Digitaria ciliaris* | Philippines | | x | - |
| | US0043 | *Digitaria sp.* | USA, Delaware, Wilmington | | x | - |
| | DsLIZ | *Digitaria sanguinalis* | USA | x | | |
| | VO107 | *Digitaria sanguinalis* | USA | x | | |
| | NI907 | *Digitaria sanguinalis* | Japan | x | | |
| | CR0027 | *Digitaria sanguinalis* | Reoublic of Korea, Jeonbug, Jeongup | | x | - |
| *Pyricularia penniseticola* | BF0017 | *Pennisetum typhoides* | Burkina Faso, Kamboinse | | x | - |

| | | | | | | |
|---|---|---|---|---|---|---|
| | CD0086 | *Pennisetum typhoides* | Côte d'Ivoire, Ferkesedougou | | x | - |
| | CD0180 | *Pennisetum sp.* | Côte d'Ivoire, Madiani | | x | - |
| | ML0031 | *Pennisetum sp.* | Mali, Sikasso | | x | - |
| | ML0048 | *Digitaria exilis* | Mali, Segou | | x | - |
| | PM1 | *Pennisetum americanum* | USA | x | | |
| *Pyricularia pennisetigena* | BR0030 | *Cenchrus echinatus* | Brazil, Goias, Goiana | | x | - |
| | BR0063 | *Cenchrus echinatus* | Brazil, Piaui | | x | - |
| | BR0067 | *Cenchrus echinatus* | Brazil, Marnhao, Imperatriz | | x | - |
| | BR0093 | *Echinochloa colona* | Brazil, Paran, Primero de maio | | x | - |
| | ML0036 | *Pennisetum sp.* | Mali, Cinzana | | x | - |
| | MX0001 | *Cenchrus ciliaris* | Mexico, Piedras negras | | x | - |
| | PH0047 | *Cenchrus echinatus* | Philippines | | x | - |
| | US0044 | *Pennisetum glaucum* | USA, Georgia, Tifton | | x | - |
| | US0045 | *Pennisetum glaucum* | USA, Georgia, Tifton | | x | - |
| | US0067 | *Cenchrus ciliaris* | USA, Texas | | x | - |
| *Pyricularia urashimae* | PJI-001 | *Oryza sativa* | China | x | | |
| | PJI-002 | *Oryza sativa* | China | x | | |
| | PJI-003 | *Oryza sativa* | China | x | | |
| | PJI-004 | *Oryza sativa* | China | x | | |
| | CBS 133598 | *Leersia oryzoides* | Japan | x | | |
| *Pyricularia zingibericola* | RN0001 | *Zingiber officinale* | France, Reunion island | | x | - |
| **Other species** | | | | | | |
| *Proxipyricularia zingiberis* | JP0041 | *Zingiber officinale* | Japan | x | x | - |
| *Pseudopyricularia cyperi* | PH0053 | *Cyperus rotundus* | Phillippines | | x | - |
| | IS0001 | *Cyperus rotondus* | Israel | | x | - |
| *Pseudopyricularia killingae* | PH0054 | *Cyperus brevifolius* | Phillippines | | x | - |

| | | | | | | |
|---|---|---|---|---|---|---|
| *Pseudopyricularia sp.* | EG0028 | *Cyperus rotundus* | Egypt | | x | - |
| | PH0052 | *Cyperus rotundus* | Philippines | | x | - |
| *Curvularia sp.* | FR9023 | *Lolium perenne* | France, Montpellier | | x | - |
| | FR9024 | *Lolium perenne* | France, Montpellier | | x | - |
| | FR9025 | *Lolium perenne* | France, Montpellier | | x | - |
| | FR9026 | *Lolium perenne* | France, Montpellier | | x | - |
| | FR9027 | *Lolium perenne* | France, Montpellier | | x | - |
| | FR9028 | *Lolium perenne* | France, Montpellier | | x | - |
| | FR9022 | *Lolium perenne* | France, Montpellier | | x | - |

**Table S2. Quantification results of *P. oryzae* obtained from the different ratios tested of infected grass in healthy grass. ND : Not detected**

| Ratio tested | Results Copies.g-1 | Mean Copies.g-1 | Standard deviation Copies.g-1 |
|---|---|---|---|
| Replicate 1 (.0.1%) | $6.00 \times 10^4$ | | |
| Replicate 2 (.0.1%) | $1.35 \times 10^4$ | $3.12 \times 10^4$ | $2.52 \times 10^4$ |
| Replicate 3 (.0.1%) | $2.02 \times 10^4$ | | |
| Replicate 1 (.0.05%) | $1.00 \times 10^4$ | | |
| Replicate 2 (.0.05%) | $2.42 \times 10^4$ | $1.30 \times 10^4$ | $1.01 \times 10^4$ |
| Replicate 3 (.0.05%) | $4.66 \times 10^3$ | | |
| Replicate 1 (.0.025%) | $1.84 \times 10^3$ | | |
| Replicate 2 (.0.025%) | ND | $9.97 \times 10^2$ | $9.28 \times 10^2$ |
| Replicate 3 (.0.025%) | $1.15 \times 10^3$ | | |
| Positive sample used (100%) | $1.47 \times 10^7$ | | |

**Table S3. Quantification of *P. oryzae* DNA (in copies.$g^{-1}$) in 3 matrices (soil, roots, and aerial parts of turf) in 15 stadiums .** Values below the detection threshold were scored 0.

| Stadium | Soil | Roots | Turf |
|---|---|---|---|
| 1 | 0 | $5.09 \times 10^4$ | 0 |
| 2 | $5.20 \times 10^2$ | $1.06 \times 10^5$ | $6.77 \times 10^4$ |
| 3 | $1.58 \times 10^3$ | $2.95 \times 10^3$ | 0 |
| 4 | 0 | 0 | 0 |
| 5 | $5.24 \times 10^2$ | $6.62 \times 10^5$ | $1.20 \times 10^3$ |
| 6 | 0 | $4.96 \times 10^4$ | 0 |
| 7 | $9.81 \times 10^2$ | 0 | 0 |
| 8 | $1.08 \times 10^3$ | 0 | 0 |
| 9 | $4.08 \times 10^3$ | $2.53 \times 10^4$ | $1.92 \times 10^3$ |
| 10 | $5.46 \times 10^2$ | $5.14 \times 10^4$ | 0 |
| 11 | $3.03 \times 10^3$ | 0 | 0 |
| 12 | $3.31 \times 10^3$ | $3.91 \times 10^5$ | $4.61 \times 10^3$ |
| 13 | 0 | $2.58 \times 10^4$ | $1.69 \times 10^4$ |
| 14 | 0 | $2.55 \times 10^4$ | 0 |
| 15 | 0 | 0 | 0 |
| Mean | $1.04 \times 10^3$ | $9.27 \times 10^4$ | $6.16 \times 10^3$ |
| Standard deviation | $1.36 \times 10^3$ | $1.86 \times 10^5$ | $1.76 \times 10^4$ |
